# Crosstalk with the GAR-3 receptor contributes to feeding defects in *Caenorhabditis elegans eat-2* mutants

**DOI:** 10.1101/562041

**Authors:** Alena A. Kozlova, Michele Lotfi, Peter G. Okkema

**Affiliations:** Department of Biological Sciences, University of Illinois at Chicago, Chicago, Illinois

**Keywords:** *C. elegans*, pharynx, GCaMP3, nicotinic acetylcholine receptor, muscarinic acetylcholine receptor, peristalsis, lifespan, feeding

## Abstract

Precise signaling at the neuromuscular junction (NMJ) is essential for proper muscle contraction. In the *C. elegans* pharynx, acetylcholine (ACh) released from the MC and M4 motor neurons stimulates two different types of contractions in adjacent muscle cells, termed pumping and isthmus peristalsis. MC stimulates rapid pumping through the nicotinic ACh receptor EAT-2, which is tightly localized at the MC NMJ, and *eat-2* mutants exhibit a slow pump rate. Surprisingly, we found that *eat-2* mutants also hyperstimulated peristaltic contractions, and these are characterized by increased and prolonged Ca^2+^ transients in the isthmus muscles. This hyperstimulation depends on crosstalk with the GAR-3 muscarinic acetylcholine receptor as *gar-3* mutation specifically suppressed the prolonged contraction and increased Ca^2+^ observed in *eat-2* mutant peristalses. Similar GAR-3 dependent hyperstimulation was also observed in mutants lacking the *ace-3* acetylcholinesterase, and we suggest that NMJ defects in *eat-2* and *ace-3* mutants result in ACh stimulation of extrasynaptic GAR-3 receptors in isthmus muscles. *gar-3* mutation also suppressed slow larval growth and prolonged lifespan phenotypes that result from dietary restriction in *eat-2* mutants, indicating that crosstalk with the GAR-3 receptor has a long-term impact on feeding behavior and *eat-2* mutant phenotypes.

**Article Summary:** Acetylcholine stimulates different contractions in adjacent muscle cells in the *C. elegans* pharynx called pumping and peristalsis. The signaling mechanisms stimulating pumping have been characterized, but how these mechanisms affect peristalsis is unknown. Here we examined muscle contractions and Ca^2+^ transients during peristalsis in wild-type animals and acetylcholine signaling mutants. Surprisingly we found that while mutants affecting the *eat-2* nicotinic acetylcholine receptor exhibited reduced pumping, they also hyperstimulated peristalses. This hyperstimulation depends on crosstalk with the GAR-3 muscarinic acetylcholine receptor in adjacent cells, and it contributes to the well-characterized dietary restriction and extended adult lifespan observed in *eat-2* mutants.

## Introduction

Communication between motor neurons and their target muscles is crucial for proper muscle contraction and function. This communication is mediated by the neurotransmitter acetylcholine (ACh), which is released from the motor neuron and binds receptors embedded in the muscle cell membrane. In vertebrate skeletal muscle, these receptors are nicotinic acetylcholine receptors (nAChRs), which are homo-or hetero-penatmeric, ligand-gated ion channels [reviewed in (Albuquerque *et al.* 2009)]. In smooth and cardiac muscle, the receptors are typically muscarinic acetylcholine receptors (mAChRs), which are 7-pass transmembrane, G-protein coupled receptors [reviewed in (Wess 2004)]. nAChRs are fast-acting receptors that rapidly and directly depolarize muscle cells to stimulate contraction in response to ACh, while mAChRs act more slowly by activating downstream signal transduction cascades and can either stimulate or inhibit contraction. A number of inherited and acquired diseases affecting communication at the neuromuscular junction in human skeletal muscle exhibit defects in nAChRs and other proteins required for their function, including myasthenia gravis and congenital myasthenic syndromes [reviewed in (Engel *et al.* 2015; Gilhus 2016)].

We are examining the mechanisms controlling contractions of the pharyngeal muscles of *Caenorhabditis elegans*. The pharynx is a tubular organ with large muscle cells positioned around a central lumen (Figure 1 A) (Albertson and Thomson 1976). The myofilaments in these muscles are oriented radially, so that contraction opens the pharyngeal lumen and relaxation closes this lumen. During feeding, these muscles perform two distinct types of contractions, termed pumping and peristalsis [reviewed in (Avery and You 2012)]. Pumping occurs frequently (~200/min), and it is a simultaneous contraction of muscles in the corpus and anterior isthmus that ingests bacteria into the lumen and concentrates this material in the anterior region of the isthmus, while at the same time contraction of muscles in the terminal bulb crushes bacteria and expels the debris into the intestine (Figure 1 B). Peristalsis occurs relatively infrequently and always following a pump. Peristalsis is a wave-like contraction followed by rapid relaxation traveling through the pm5 muscle that transports a bolus of ingested bacteria from the anterior isthmus to the terminal bulb (Figure 1 C). The contractions in the pm5 muscle are remarkably complex. Three pm5 muscles are arranged around the circumference of the pharyngeal lumen, and each of these cells extends through the entire length of the isthmus (Figure 1 A) (Albertson and Thomson 1976). During a pump, the anterior portion of the pm5 cells contracts to open the lumen in the anterior isthmus, but the posterior half of these cells remain relaxed. In contrast, during peristalsis, the anterior portion of pm5 remains relaxed, while a wave-like contraction followed by relaxation travels from the center to posterior isthmus.

**Figure 1:**
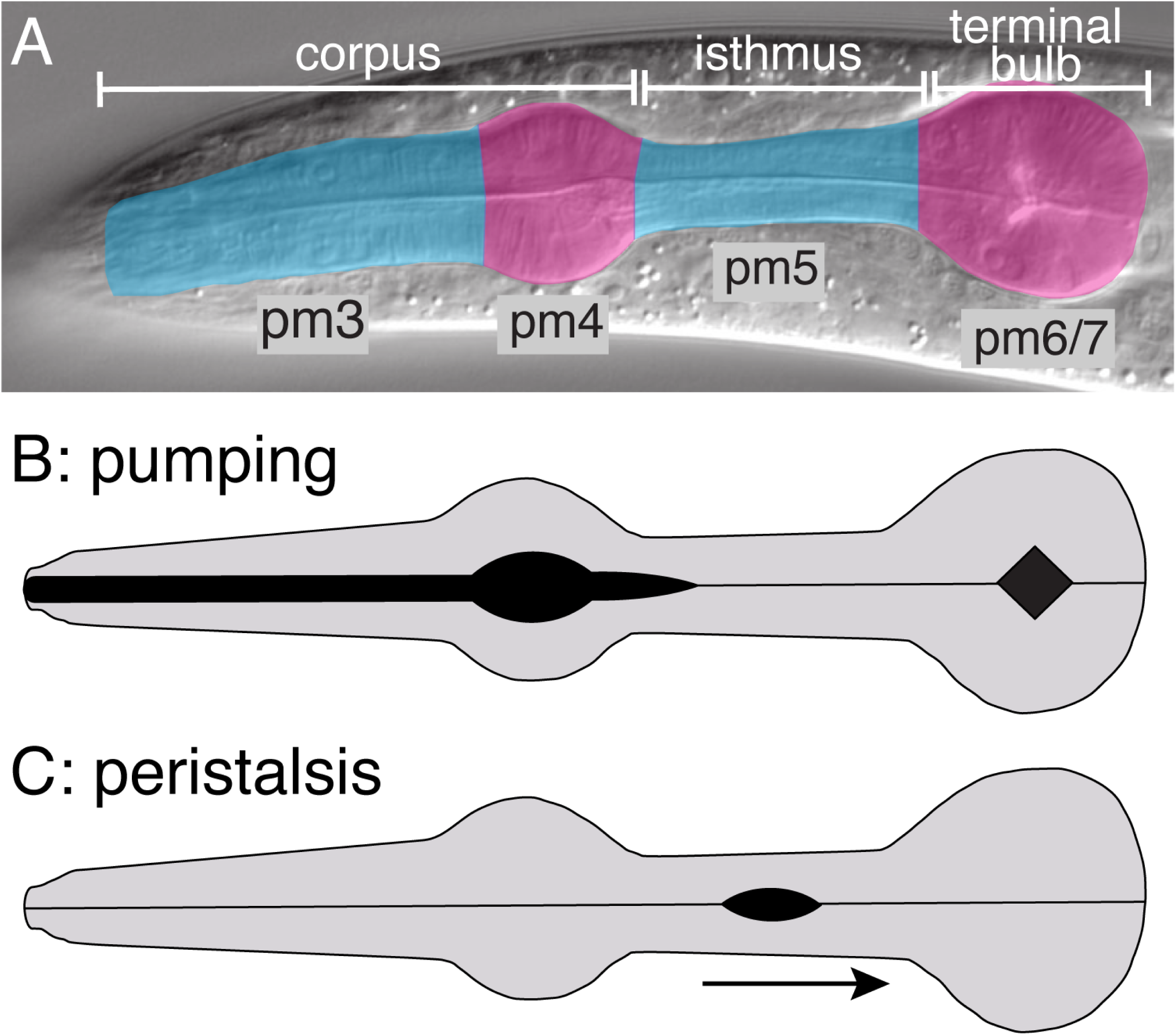
Pharyngeal anatomy and contractions. (A) DIC micrograph of an adult pharynx indicating anatomical regions (white bars) and colored to indicate the location of pharyngeal muscle cells (pm). The pharynx exhibits 3-fold rotational symmetry, and there are three of the pm3-pm7 muscle cells surrounding the central lumen (Albertson, 1976). pm5 cells extend the length of the isthmus and are the primary focus of this work. (B, C) Diagrams indicating pharyngeal muscle contractions during pumping and peristalsis. The black regions indicate open lumen, and the arrow indicates the direction of the peristaltic contraction.

Rapid pumping is stimulated by the two MC neurons that form synapses on the muscles near the junction of the pm4 and pm5 muscles (Mckay *et al.* 2004), and laser ablation of the MCs results in a slow pump rate (Avery and Horvitz 1989). The MCs stimulate a nAChR containing the non-alpha subunit EAT-2 that clusters at the MC synapses, and *eat-2* mutants exhibit the same slow pump rate observed in MC ablated animals (Raizen and Avery 1994; Mckay *et al.* 2004). EAT-2 stimulation initiates an increase in cytoplasmic Ca^2+^ concentration in the pharyngeal muscles that is mediated by the voltage-activated Ca^2+^ channels CCA-1 and EGL-19 leading to muscle contraction (Shtonda and Avery 2005; Steger *et al.* 2005; Kerr 2006). MC does not synapse on the terminal bulb, but simultaneous contraction of corpus and terminal bulb is mediated by electrical coupling (Starich *et al.* 1996). In the absence of MC or EAT-2, pumping occurs slowly, and it is believed that these contractions are generated autonomously within the muscle.

Peristalsis is stimulated by the single M4 motor neuron that extends processes through the isthmus and synapses on the muscles in the posterior half of pm5 (Albertson and Thomson 1976), and ablation of M4 eliminates peristalsis (Avery and Horvitz 1987). The receptors that respond to M4 to produce a peristalsis are unknown, but our previous studies of M4 defective mutants have implicated mAChRs (Ray *et al.* 2008; Ramakrishnan and Okkema 2014), while other studies suggest that peptide neurotransmitters secreted from other neurons may also be involved (Song and Avery 2012). The *C. elegans* genome contains three genes encoding mAChRs, *gar-1, gar-2,* and *gar-3*, but only *gar-3* is expressed in pharyngeal muscles (Lee *et al.* 2000; Steger and Avery 2004; Dittman and Kaplan 2008), and the muscarinic agonist arecoline can stimulate pumping through GAR-3 (Steger and Avery 2004).

Here we examine the roles nAChRs and mAChRs play in peristalsis in wild-type animals and mutants defective in ACh synthesis or response, using both direct observation and imaging the genetically encoded Ca^2+^ indicator GCaMP3 (Tian *et al.* 2009). We find that mutants lacking endogenous ACh fail to pump or peristalse indicating that pharyngeal muscle contractions are not generated autonomously. Further, treatment of these mutants with either nicotinic or muscarinic agonists stimulated both pumping and peristaltic contractions, and the response to these agonists depends on the nAChR subunit EAT-2 and the mAChR GAR-3, respectively. Surprisingly, in the absence of exogenous agonist, *eat-2* mutants exhibited hyperstimulated isthmus muscle peristalses and increased isthmus muscle Ca^2+^ concentrations, and that this hyperstimulation depends on crosstalk with the GAR-3 receptor. This relationship between EAT-2 and GAR-3 affected feeding behavior throughout the life of the animal, as *gar-3* mutation suppressed the slow larval growth and prolonged lifespan phenotypes resulting from dietary restriction in *eat-2* mutants (Avery 1993a; Lakowski and Hekimi 1998). *ace-3* acetylcholinesterase mutants similarly exhibited hyperstimulated isthmus peristalses that were dependent on *gar-3*, and we hypothesize that in the absence of EAT-2 or ACE-3 unbound ACh spills over from the synapse and stimulates extrasynaptic GAR-3 in the isthmus muscles.

## Materials and Methods

### Nematode handling, transformation and strains

*C. elegans* strains were grown under standard conditions (Lewis and Fleming 1995). Germline transformations were performed by microinjection using pRF4 (100 ng/µl) containing *rol-6(su1006)* as a transformation marker and the *myo-2^Prom^::GCaMP3* reporter pOK294.04 (20 ng/µl generating *cuEx804*) or the *ceh-19b^Prom^::snb-1::gfp* reporter pOK361.01(25 ng/µl generating *cuEx826*) (Mello and Fire 1995). The extrachromosomal *myo-2^Prom^::GCaMP3* transgene *cuEx804* was chromosomally integrated by UV/TMP mutagenesis and outcrossed to form *cuIs36*.

The following strains were generated by the *C. elegans* Gene Knockout Consortium (Consortium 2012): VC1836 *cha-1(ok2253); nT1[qls51],* VC657 *gar-3(gk305),* FX863 *acr-7(tm863),* RB918 *acr-16(ok789),* VC2661 *acr-10(ok3118),* RB2294 *acr-6(ok3117),* VC2820 *eat-2(ok3528)/mT1; +/mT1*.

The following strains were constructed by others: wild-type strain N2 (Brenner 1974), RB1132 *acr-14(ok1155)* (Ruaud and Bessereau 2006), DA1116 *eat-2(ad1116)* (Raizen *et al.* 1995), DA465 *eat-2(ad465)* (Avery 1993a), PR1300 *ace-3(dc2)* (Combes *et al.* 2000), GC201 *ace-2(g72); ace-1(p1000)* (Talesa *et al.* 1995; Culetto *et al.* 1999)*, eat-18(ad820)* (Raizen *et al.* 1995), and UL2702 *unc-119(ed3); leIs2702(ceh-19b^Prom^::gfp,unc-119(+))* (Feng and Hope 2013).

The following strains were constructed in this study: OK1020 *cuIs36[myo-2^Prom^::GCaMP3]*, OK1062 *gar-3(gk305); cuIs36*, OK1063 *eat-2(ok3528); cuIs36*, OK1064 *eat-2(ok3528); gar-3(gk305),* OK1075 *eat-2(ok3528); gar-3(gk305); cuIs36,* OK1023 *eat-2(ok3528)*, OK1081 *ace-3(dc2); gar-3(gk305)*, OK1082 *eat-2(ok3528); unc-119(ed3); leIs2702(ceh-19b^Prom^::gfp, unc-119(+))*; OK1083 *cuEx828[ceh-19b^Prom^::snb-1::gfp*]; and OK1084 *eat-2(ok3528); cuEx828*.

### General methods for nucleic acid manipulations and plasmid construction

Standard methods were used to manipulate all DNAs (Ausubel 1990), and plasmids sequences are available from the authors. The *myo-2* promoter from plasmid pPD96.48 was cloned into HindIII and MscI digested *str-2::GCaMP3* plasmid (Chalasani *et al.* 2007) to generate pOK294.04. The *ceh-19b* promoter was amplified from N2 genomic DNA using primers PO1452 (CGAGCATGCGAAAAACAGGAAAGTCTCG) and PO1453 (GACCCGGGATGTAGAGTTGAGAAGTTGCCA), digested with SphI and XmaI and inserted into SphI and XmaI digested *ser-7b^Prom^::snb-1::gfp* plasmid pOK219.08 to generate the *ceh-19b^Prom^::snb-1::gfp* plasmid pOK361.01 (Ray *et al.* 2008).

### Genotyping

Individual animals were genotyped for *eat-2(ok3528)*, *gar-3(gk305)* and *ace-3(dc2)* alleles by PCR (Beaster-JONES AND OKKEMA 2004). Primers for genotyping *eat-2(ok3528)* were PO1423 (TGCGTGGTAGAGGGATAGTG), PO1424 (TCTCGACGAGACCTACGTTG), PO1425 (ACAGCTACAGTACCTCGCAC). Primers for genotyping *gar-3(gk305)* were PO1426 (TAATAGGTTCGGCCCAGAGC), PO1427 (GTGATCGTTTGCTGGGAAGC), PO1428 (CGAAGCTCAGAATGTCAGTAACG). Primers for genotyping *ace-3(dc2)* were PO1435 (CAAGGATACAGAGTACACGGCA), PO1436 (CAAGCCCGCAAATTGAACTGA), PO1437 (GCAAGTGGCAAGCGAGAATA).

### Analysis of feeding behavior and drug studies

To analyze pharyngeal muscle contractions, L1 larvae hatched in the absence of food were suspended in 5 μl of M9 buffer containing OP50 and imaged on a 2% agarose pad under a coverslip. For drug treatments, either arecoline (Acros Organics, Cat# CAS: 300-08-3) or nicotine (Sigma, Cat# N5260-25G) was included and animals were treated for 15 min prior to adding a coverslip. Individual N2 or mutant animals that pumped were recorded at 25 frames/sec for 1 min (*cha-1* mutants were recorded for 5 min) using a Zeiss AxioImager microscope with an MRm camera and ZEN Software. For each genotype and drug treatment the feeding behavior was analyzed in at least 5 animals (~500 pumps/per animal). Time-lapse images were exported and processed using Fiji/ImageJ (Schindelin *et al.* 2012) to generate time-lapse movies. Acquisition times were exported from ZEN and quantifications were performed using Microsoft Excel.

### Calcium imaging and microscopy

Young adult animals were incubated in 5 μl of 20mM serotonin (Sigma, Cat# H7752-5G) for 10 min on 2% agarose pad and then immobilized using 1.5 μl Polystyrene 0.10 micron microspheres and a coverslip (Polysciences, Inc, Cat# 00876). GCaMP3 imaging was performed on a Zeiss AxioImager microscope using a Q-Imaging Rolera EM-C^2^ EMCCD camera. Time-lapse movies were captured at 25-30 frames/sec using ZEN software and 14-bit TIFF images were exported. Animals pumping with frequency less than 100 pumps/min were analyzed. For quantification, the pharynx was straightened with CellProfiler (Kamentsky *et al.* 2011) and aligned using the StackReg Fiji plugin (Thevenaz *et al.* 1998). The isthmus was cropped and two regions of interest (ROI) were drawn as 10 pixel (2.54 µm) wide lines in the center and posterior isthmus respectively. Fluorescence measurements were analyzed using custom Matlab scripts or using Microsoft Excel. Total fluorescence was measured within the ROI and normalized for each GCaMP3 peak using formula: normalized ΔF=(Fmax-Fmin)/Fmin*100% (where Fmax is maximum fluorescence of GCaMP3 peak and Fmin is a minimum fluorescence immediately before GCaMP3 increase). Peak duration was quantified as a width of GCaMP3 fluorescence at half peak height. Rise time was quantified as time that it took for each fluorescence peak to reach its maximum. A Student's t-test was used to compare GCaMP3 fluorescence measurements between different genotypes, and boxplots were generated using the Matlab *boxplot* command. Peak delay was calculated as time between the maximum increase in GCaMP3 fluorescence in the center and posterior isthmus using the Matlab *diff* command. Average baseline fluorescence levels in the entire pharyngeal isthmus were measured in a representative image during an interpump period acquired at a time point early in time lapse image sets of each animal using Fiji/ImageJ.

GFP and DIC images captured using a Zeiss Axiocam MRm and Zeiss Zen software. To characterize *ceh-19b^Prom^::gfp* expression, Z-series images were collected. Maximum intensity Z projections were produced using Fiji/ImageJ (Schindelin *et al.* 2012). *ceh-19b^Prom^::snb-1::gfp* expression was faint, and Z-series were collected in adult animals immobilized in 10 mM NaN_3_ and false colored in Fiji/ImageJ using the Rainbow RBG LUT.

### Growth assay

Adult hermaphrodites were allowed to lay eggs for 8 hours at 25°C, and embryos were transferred to freshly seeded plates and incubated at 20°C for 16 hours. Unhatched embryos were counted, and hatched L1s were allowed to grow an additional three days. Adult animals were counted and removed on day 5. *eat-2(ok3528)* and *eat-2(ok3528); gar-3(gk305)* were allowed to grow for an additional day to quantify slow growing adults. For each genotype, two plates were set up with 50 embryos each.

### Life span assay

Life span assays were performed according to previously published protocol (Sutphin and Kaeberlein 2009). Assays were performed starting with 30 L4 worms fed with UV-killed OP50 *E. coli* at 20°C, and dead or missing animals were replaced by animals of the same age. For each experiment, triplicate plates of each genotype were assayed, and representative results of 3 or more experiment are shown. Kaplan Meier Survival Analysis was used to generate survival plots and calculate median life span, and for statistical analysis (GraphPad Prism 5).

## Results

### Pumping and isthmus peristalsis can be stimulated through both nicotinic and muscarinic receptors

We are interested in the signaling mechanisms that produce pharyngeal muscle contractions and productive feeding. M4 and the MCs are cholinergic motor neurons that express choline acetyltransferase encoded by *cha-1* (Raizen *et al.* 1995; Ramakrishnan and Okkema 2014; Pereira *et al.* 2015), and strong *cha-1* mutants have reduced pharyngeal contractions and arrest as severely uncoordinated L1s (Rand 1989; Avery and Horvitz 1990). Previous studies have shown that stimulation of nicotinic acetylcholine receptors (nAChRs) with nicotine or muscarinic acetylcholine receptors (mAChRs) with arecoline could induce pharyngeal muscle contractions, but these studies did not specifically examine peristalses in the absence of endogenous acetylcholine (Avery and Horvitz 1990; Raizen *et al.* 1995; Ramakrishnan and Okkema 2014).

To extend this work, we examined the effect of nicotine or arecoline treatment on *cha-1(ok2253)* mutant L1 animals. *cha-1(ok2253)* is a previously uncharacterized null allele containing a 1.7 kb deletion that eliminates the catalytic histidine (His341) required for ACh synthesis. and homozygous mutants arrest as paralyzed L1s with a coiled appearance (Consortium 2012). Untreated *cha-1(ok2253)* homozygotes completely lacked pharyngeal muscle contractions, but mutant animals treated with either nicotine or arecoline exhibited both pumping and peristalsis (Figure 2; Movie S1-S4). These results demonstrate that acetylcholine is necessary for pumping and peristalsis indicating that pharyngeal muscles do not have autonomous contractile activity. Further, activation of either nAChRs or mAChRs can stimulate these contractions. Notably we observed occasional animals treated with either agonist that exhibited peristaltic contractions in the isthmus without a preceding pump (2/10 animals treated with 10 mM arecoline; 2/40 animals treated with 5 mM nicotine), consistent with our previous observations that peristalsis can be uncoupled from pumping by directly stimulating receptors in the isthmus muscles (Ramakrishnan and Okkema 2014). In addition, while higher arecoline concentrations increased the percentage of animals exhibiting pharyngeal muscle contractions, higher nicotine concentration progressively decreased the percentage of animals exhibiting pharyngeal muscle contractions. This decrease may result from hyperstimulation of nAChRs as has been previously reported (Avery and Horvitz 1990), although we did not observe the tetanic contractions previously observed in animals treated with nicotine.

**Figure 2:**
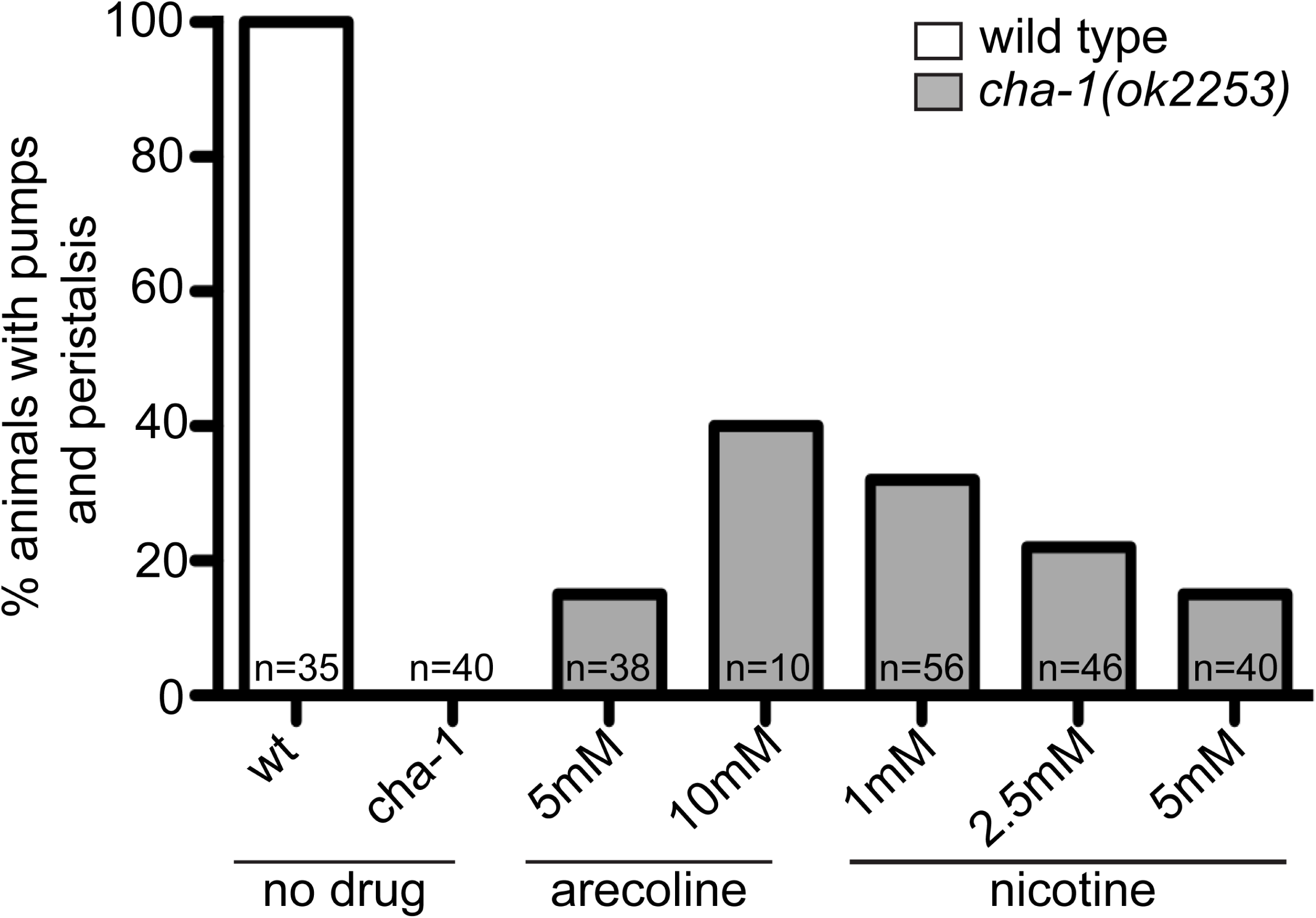
Muscarinic and nicotinic receptor agonists stimulate pharyngeal muscle contractions in *cha-1* mutants. The percentage of wild-type or *cha-1(ok2253)* L1 animals exhibiting pharyngeal muscle pumping and peristalses, either untreated or treated with the indicated concentrations of arecoline or nicotine. Animals were visualized for 5 min each, and the number of animals observed (n) is indicated.

### EAT-2 and GAR-3 mediate the response to agonists

The results above indicate that activation of either mAChRs or nAChRs can stimulate pumping and peristaltic contractions in the pharynx. To identify the receptors involved in these responses, we asked if mutants affecting receptors expressed in the pharyngeal muscles become insensitive to agonists.

To examine the role mAChR signaling plays in pharyngeal muscle contractions, we characterized these contractions in *gar-3(gk305)* null mutants (Liu *et al.* 2007). *gar-3(gk305)* mutants are viable, look healthy, and grow normally, and pharyngeal muscle contractions in *gar-3(gk305)* L1s were similar to those in wild type (Table 1). Neither the rate of pumping nor the frequency or duration of peristalsis was significantly different between wild-type and *gar-3(gk305)* animals. In comparison, *gar-3(gk305)* animals were almost completely insensitive to exogenously applied arecoline (Table 1). Wild-type animals treated with arecoline exhibited a strongly reduced pump rate, and each of these pumps was followed by a peristalses that was prolonged compared to untreated animals. In contrast, *gar-3(gk305)* mutants were largely unaffected by arecoline. The pump rate was unaffected in *gar-3(gk305)*, and, although these animals did exhibit a small decrease in the duration of peristaltic contractions when treated with arecoline, we note that this change is the opposite of the prolonged peristalses observed in arecoline treated wild-type animals. Thus, GAR-3 mediates the response to arecoline, but loss of this receptor has no strong effect on pharyngeal muscle contractions in the absence of this drug.

**Table 1:** Quantification of pharyngeal muscle contractions

| genotype (treatment) | pump rate<br>(pumps/min ± sem) | peristalsis rate<br>(peristalsis/min ± sem) | % pumps followed by peristalsis | duration of isthmus peristalsis<br>(ms ± sem) |
| --- | --- | --- | --- | --- |
| wild type <sup>a</sup> | 218 ± 11 | 31 ± 3 | 14% | 113 ± 4 |
| wild type + arecoline <sup>b</sup> | 13 ± 2 | 13 ± 2 | 100% | 696 ± 92 |
| *gar-3(gk305)* <sup>c</sup> | 183 ± 15 | 32 ± 6 | 18% | 146 ± 6 |
| *gar-3(gk305)* + arecoline <sup>d</sup> | 173 ± 13 | 18 ± 4 | 10% | 119 ± 5 |
| *eat-2(ad465)* <sup>e</sup> | 66 ± 17 | 39 ± 11 | 59% | 236 ± 11 |
| *eat-2(ad1116)* <sup>f</sup> | 32 ± 5 | 18 ± 2 | 63% | 246 ± 9 |
| *eat-2(ok3528)* <sup>g</sup> | 27 ± 4 | 24 ± 3 | 86% | 398 ± 12 |
| *eat-2(ok3528); gar-3(gk305)* <sup>h</sup> | 20 ± 2 | 15 ± 3 | 75% | 227 ± 7 |
| *eat-18(ad820)* <sup>i</sup> | 19 ± 3 | 18 ± 3 | 92% | 378 ± 26 |
| *ace-2(g72); ace-1(p1000)* <sup>j</sup> | 132 ± 38 | 20 ± 6 | 16% | 191 ± 12 |
| *ace-3(dc2)* <sup>k</sup> | 128 ± 30 | 14 ± 2 | 14% | 305 ± 14 |
| *ace-3(dc2); gar-3(gk305)* <sup>l</sup> | 235 ± 20 | 36 ± 7 | 15% | 82 ± 1 |

<sup>a</sup> 17 N2 L1s were recorded for 19-20 s and a total of 1194 pumps were analyzed.
<sup>b</sup> Six N2 L1s treated with 5 mM arecoline were recorded for 34-46 s and a total of 48 pumps were analyzed. These animals exhibited a reduced pump rate and prolonged peristalses compared to untreated animals ( $p < 10^{-5}$ ).
<sup>c</sup> Six *gar-3(gk305)* L1s were recorded for 19-21 s and a total of 361 pumps were analyzed.
<sup>d</sup> Six *gar-3(gk305)* L1s treated with 5 mM arecoline were recorded for 19-21 s and a total of 342 pumps were analyzed. The pump rate was unaffected, but the duration of isthmus peristalsis was decreased compared to untreated *gar-3(gk305)* mutants ( $p < 10^{-4}$ ).
<sup>e</sup> Five *eat-2(ad465)* L1s were recorded for 21-22 s and a total of 75 pumps were analyzed. The pump rate was decreased ( $p < 10^{-4}$ ) and the duration of isthmus peristalsis was increased ( $p < 10^{-21}$ ) compared to wild type.
<sup>f</sup> Five *eat-2(ad1116)* L1s were recorded for 20-21 s and a total of 65 pumps were analyzed. The pump rate was decreased ( $p < 10^{-5}$ ) and the duration of isthmus peristalsis was increased ( $p < 10^{-20}$ ) compared to wild type.
<sup>g</sup> Twelve *eat-2(ok3528)* L1s were recorded for 24-25 s and a total of 121 pumps were analyzed. The pump rate was decreased ( $p < 10^{-5}$ ) and the duration of isthmus peristalsis was increased ( $p < 10^{-20}$ ) compared to wild type.
<sup>h</sup> Seven *eat-2; gar-3* L1s were recorded for 32-33 s and a total of 75 pumps were analyzed. The pump rate was unaffected, but the duration of isthmus peristalsis was reduced ( $p < 10^{-24}$ ) compared to *eat-2(ok3528)* single mutants.
<sup>i</sup> Six *eat-18* L1s were recorded for 32-33 s and a total of 59 pumps were analyzed. The duration of isthmus peristalsis was increased ( $p < 10^{-9}$ ) compared to wild type.
<sup>j</sup> Five *ace-2; ace-1* L1s were recorded for 23-24 s and a total of 258 pumps were analyzed. The duration of isthmus peristalsis was increased ( $p < 0.02$ ) compared to wild type.
<sup>k</sup> Five *ace-3* L1s were recorded for 20-21 s and a total of 226 pumps were analyzed. The duration of isthmus peristalsis was increased ( $p < 10^{-9}$ ) compared to wild type.
<sup>l</sup> Six *ace-3; gar-3* L1s were recorded for 19-20 s and a total of 468 pumps were analyzed.

We then focused on *eat-2*, which encodes the only nAChR subunit known to function in the pharyngeal muscles (Raizen *et al.* 1995; Mckay *et al.* 2004). To test if *eat-2* is necessary to respond to nicotine, we compared wild-type animals and *eat-2(ok3528)* mutants treated with nicotine at the L1 stage. Like *cha-1* mutants, wild-type animals treated with increasing concentrations of nicotine exhibited a dose-dependent decrease in pharyngeal muscle contractions (Figure 3), but *eat-2* animals were not significantly affected by this treatment. Thus the pharyngeal muscle response to nicotine depends on EAT-2 containing nAChRs.

**Figure 3:**
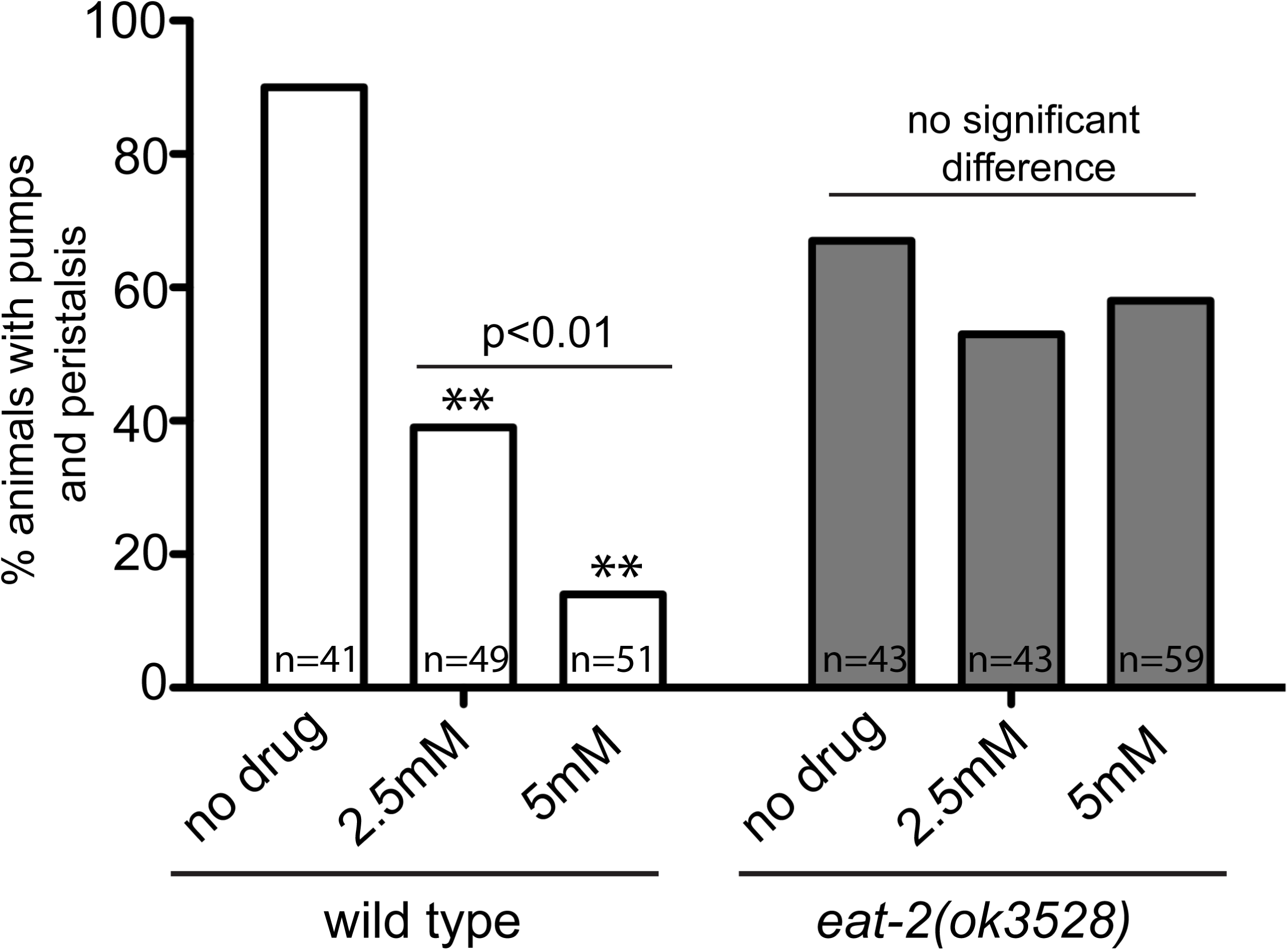
*eat-2* mutants are insensitive to exogenous nicotine. Percentage of wild-type or *eat-2(ok3528)* L1 animals exhibiting pharyngeal muscle pumping and peristalses, either untreated or treated with the indicated concentrations of nicotine. ** indicates significantly different than untreated animals (p < 0.0001), and the bar indicates significant difference between wild-type animals treated with increasing concentrations of nicotine. Animals were visualized for 5 min each, and the number of animals observed (n) is indicated.

### *eat-2* mutants exhibit prolonged pumps and peristalses

To understand how EAT-2 affects muscle contractions, we examined three strong *eat-2* mutants at the L1 stage. *eat-2(ad465)* and *eat-2(ad1116)* are point mutations introducing an early stop codon and affecting a splice site, respectively (Mckay *et al.* 2004)(WormBase WBVar00000089), while *eat-2(ok3528)* contains a 614 bp deletion predicted to cause a frameshift mutation upstream of the transmembrane domains (Consortium 2012). As expected, the rate of pumping in all three mutants was significantly reduced compared to wild type (Table 1) (Avery 1993a; Raizen *et al.* 1995). However, the duration of peristaltic contractions in the posterior isthmus was unexpectedly increased up to nearly 4-fold, with relaxation of the posterior region of the isthmus muscles particularly delayed (Movie S5). The percent of pumps that were followed by a peristalsis was also increased in *eat-2* mutants, but as the pump rate was decreased in these animals, the absolute rate of peristalsis was similar in wild-type animals and *eat-2* mutants. Notably, *eat-2(ok3528)* mutants exhibited the strongest phenotypes, which, together with molecular lesion in this allele, suggests *eat-2(ok3528)* is a null allele.

We next examined MC in *eat-2(ok3528)* mutants using a *ceh-19b^Prom^::gfp* reporter (Feng and Hope 2013) and found that MC morphology is similar in wild-type and *eat-2(ok3528)* adults (Figure 4). The MC cell bodies were located in the anterior bulb of the pharynx, and these cells extended processes posteriorly into the pharyngeal isthmus. Varicosities in these processes that mark synaptic sites were observed in both strains at the junction between the pm4 and pm5 muscles and in the anterior region of the isthmus. We then looked specifically at the locations of synapses in MC using a *snb-1::gfp* fusion expressed using the *ceh-19b* promoter. SNB-1::GFP is a functional synaptobrevin that marks synaptic vesicle clusters in presynaptic cells (Nonet 1999). In both wild-type adults and *eat-2(ok3528)* mutants, synapses were visible in the MC process beginning near the junction of the pm4 and pm5 muscle cells and extending into the anterior region of the isthmus (Figure 5). *eat-2(ok3528)* mutants exhibited a small but not statistically significant decrease in the number of vesicle clusters compared to wild type (wild type 7.0±2.40 clusters, n=8; *eat-2(ok3528)* 4.9± 1.0; p=0.09, n=13). Thus, the prolonged peristalses in *eat-2* mutants do not result from morphological or synaptic defects in the MCs. Notably using both of these reporters, we observed the MC axon extending into the anterior isthmus further than has been previously reported (Albertson and Thomson 1976) and forming synapses directly on the pm5 muscles.

**Figure 4:**
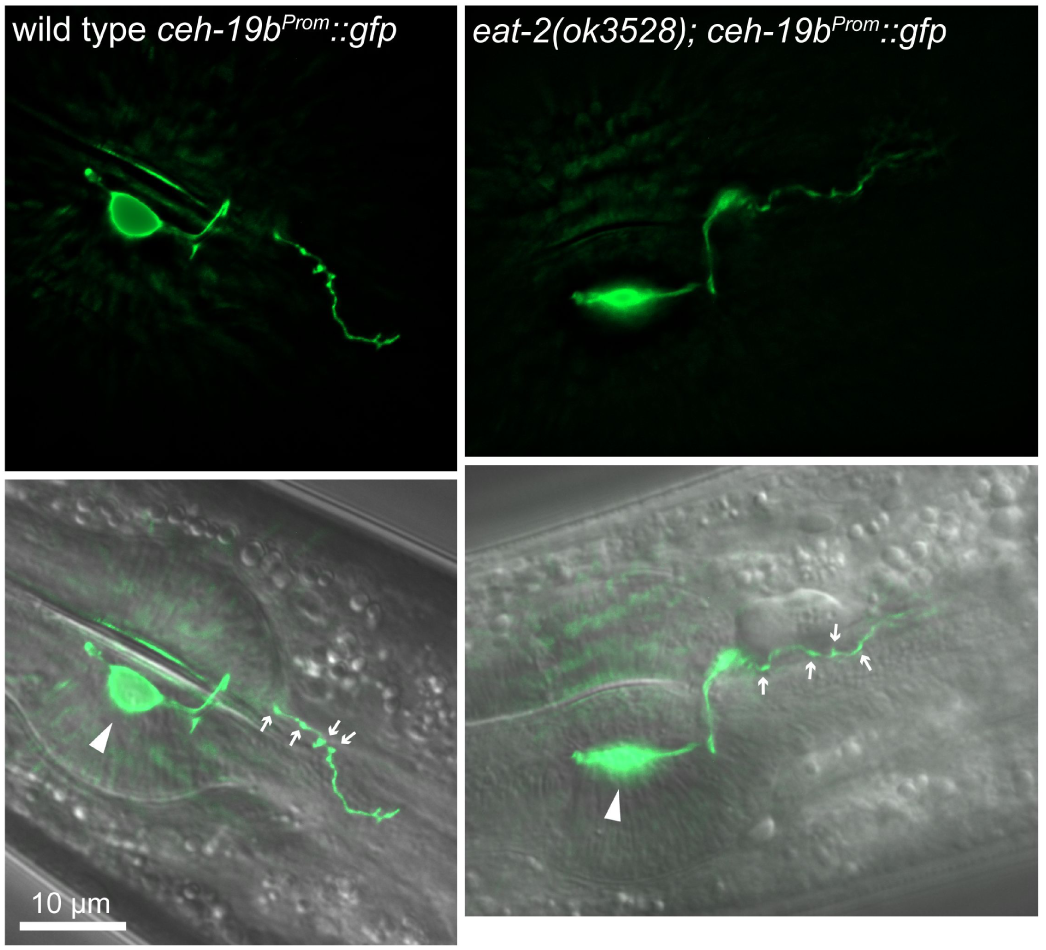
MC neuron morphology in wild-type and *eat-2* mutant adults. Fluorescence (top) and merged fluorescence & DIC images of adult animals of the indicated genotypes expressing *ceh-19b^Prom^::gfp* in the MC neuron. Anterior is left, and the metacorpus and anterior isthmus are shown. The MC cell body (arrowhead) and varicosities in the MC process (small arrows) are indicated. Fluorescence images are maximal intensity Z-projections of images through one MC cell and process.

**Figure 5:**
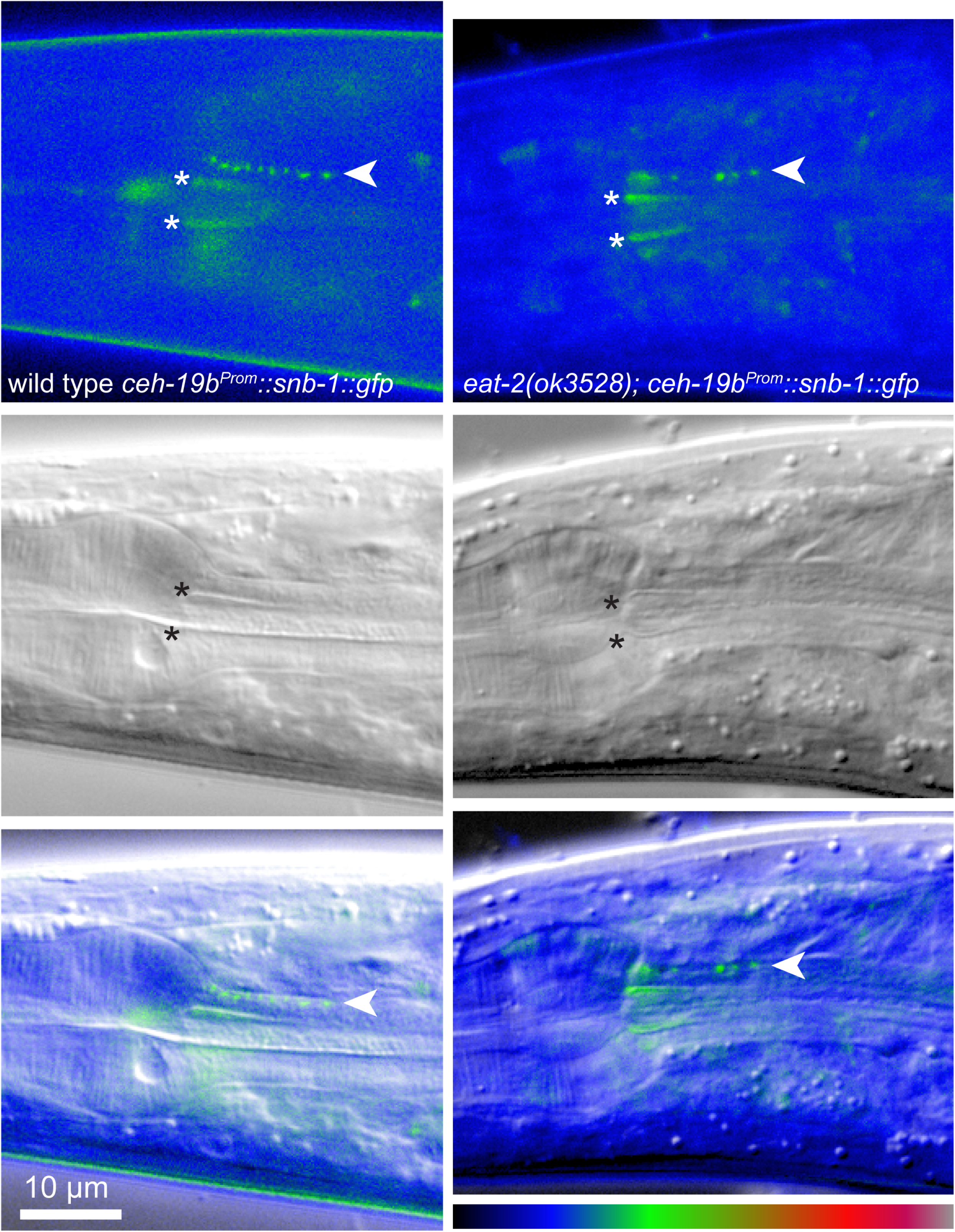
MC neuron synapses in wild-type and *eat-2* mutant adults. Fluorescence (top), DIC (middle), and merged images (bottom) of the adult animals of the indicated genotypes expressing *ceh-19b^Prom^::snb-1::gfp* in the MC neuron. Anterior is left, and the metacorpus and anterior isthmus are shown. Synaptic vesicle clusters marking *en passant* synapses in the axon of one MC neuron are marked in the fluorescence and merged images (arrowheads), Non-specific autofluorescence at the edges of the pharyngeal lumen are marked in the fluorescence and DIC images (asterisks). Fluorescence images are false colored using the look-up table at the bottom right.

To corroborate our observations in *eat-2* mutants, we also examined pharyngeal muscle contractions in *eat-18* mutants. *eat-18* encodes a novel transmembrane protein required for function of EAT-2 and other nAChRs in the pharynx, and *eat-18* mutants share the reduced pump rate with *eat-2* mutants (Raizen *et al.* 1995; Mckay *et al.* 2004). We found *eat-18(ad820)* animals also exhibit prolonged peristalses with almost all pumps followed by peristalses (Table 1). In contrast, mutants affecting several other nAChR subunits reported to be expressed in the pharyngeal muscles, including *acr-6(ok3117), acr-7(tm863), acr-10(ok3118), acr-14(ok1155)*, and *acr-16(ok789)* (Saur *et al.* 2013), did not exhibit significant changes in pumping frequency, peristalsis frequency, or the duration of peristalsis (Table S1).

Taken together these results indicate that loss of the EAT-2 nAChR subunit results in slow pumping and prolonged peristalses that occur after nearly every pump. Paradoxically the *eat-2* mutant defects in peristalsis are opposite those in pumping, indicating that, while wild-type EAT-2 stimulates rapid pumping, it also limits the duration of peristaltic contractions in the isthmus muscles.

### *gar-3* mutation suppresses the peristalsis defects in *eat-2* mutants

Pharyngeal muscle contractions in *eat-2* mutants are strikingly similar to those of wild-type animals treated with arecoline (Table 1), suggesting that some of the *eat-2* mutant phenotypes are related to mAChR signaling. Since GAR-3 is the mAChR responding to arecoline in the pharynx, we examined pharyngeal muscle contractions in *eat-2(ok3528); gar-3(gk305)* double mutants.

We found that *gar-3* mutation partially suppressed the prolonged peristalses observed in *eat-2* single mutants (Table 1). The duration of peristalses in *eat-2(ok3528); gar-3(gk305)* double mutants was strongly reduced compared to *eat-2(ok3528)* single mutants, although they were still longer than those in wild-type animals or *gar-3(gk305)* single mutants. In contrast, the reduced pumping frequency and increased frequency of peristalsis observed in *eat-2(ok3528)* single mutants was not suppressed in *eat-2(ok3528); gar-3(gk305)*. Thus, *eat-2* loss-of-function affects pumping and peristalsis by different mechanisms, and only the prolonged peristalsis phenotype depends on crosstalk with the GAR-3 receptor.

### Ca^2+^ transients in the pharyngeal isthmus parallel the contractile phenotypes in *eat-2* and *gar-3* mutants

To characterize isthmus muscle excitation in wild-type animals and mutants, we constructed strains expressing the genetically encoded Ca^2+^ indicator (GECI) GCaMP3 using the pharyngeal muscle-specific *myo-2* promoter (Okkema *et al.* 1993; Tian *et al.* 2009). GCaMP3 fluorescence increases in muscle cells as cytoplasmic Ca^2+^ concentrations increase during excitation contraction coupling [reviewed (Bers 2002)]. Similar to previous analyses of GECIs in the pharynx, we characterized adult animals treated with serotonin to stimulate pharyngeal muscle contractions and focused on animals pumping slowly to resolve individual excitation events (<100 pumps/min; <1.67 Hz) (Shimozono *et al.* 2004; Kerr 2006). Baseline fluorescence levels in the isthmus muscles were similar in the strains examined (Figure S1), and representative traces of time-lapse fluorescence are shown in Figure S2.

We observed changes in Ca^2+^ concentration in the isthmus muscles that were very dynamic (Figure 6; Movie S6). As reported in previous studies, we observed that wild-type animals displayed an increase in Ca^2+^ throughout the central isthmus during pumping followed by a longer and delayed increase in Ca^2+^ in the posterior isthmus during peristalsis (Figure 7 A, B; Movie S6) (Shimozono *et al.* 2004). We found that this Ca^2+^ signal during peristalsis exhibited a wave-like increase and decrease that traveled in an anterior to posterior direction through the posterior isthmus and resembled the progression of open pharyngeal lumen during peristalsis (Movie S7). While Ca^2+^ waves have not been previously described in the *C. elegans* pharynx (Shimozono *et al.* 2004), similar waves of Ca^2+^ can be observed in individual cardiac muscle cells, which are generated by progressive release of Ca^2+^ released from adjacent sites in the sarcoplasmic reticulum [reviewed in (Stuyvers *et al.* 2000)]. This mechanism may also underlie the wave-like contraction of pm5 during peristalsis.

**Figure 6:**
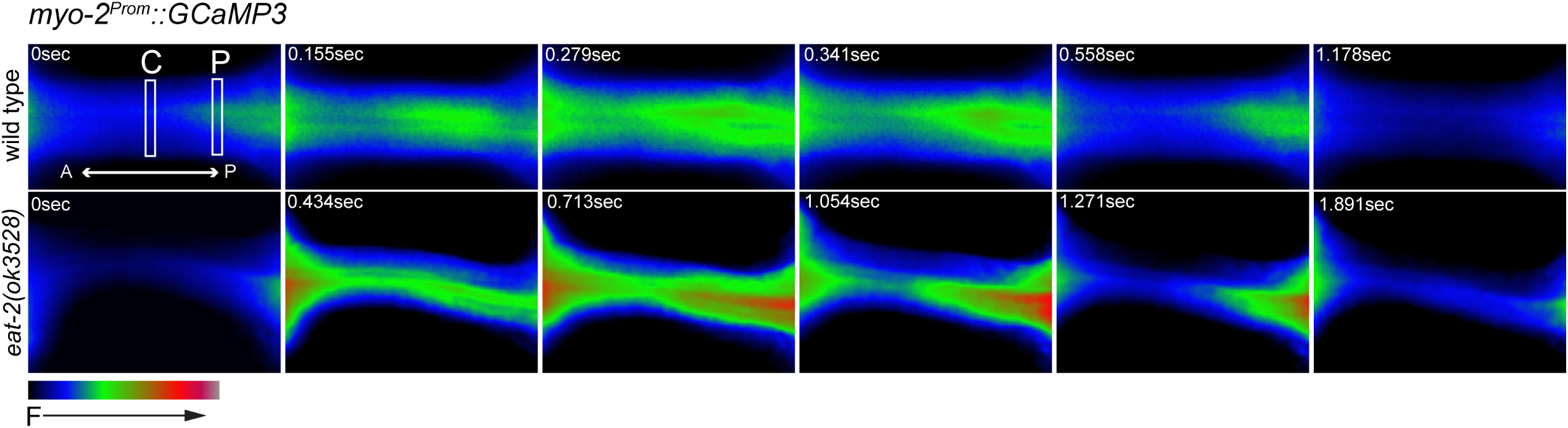
Dynamic changes in Ca^2+^ levels in the isthmus muscles. Time-lapse fluorescence images of the pharyngeal isthmus of a wild-type (top) or *eat-2(ok3528)* (bottom) adults expressing the GECI GCaMP3 in the pharyngeal muscles. Images are false colored as indicated at the lower left, and one pump and peristalsis are shown. Boxes indicate the regions where fluorescence levels were quantified in the center (C) and posterior (P) isthmus. The amount of time after fluorescence begins increasing are indicated, and the frames indicating maximum fluorescence in wild-type and *eat-2(ok3528)* are shown at 0.279 sec and 0.713 sec, respectively.

**Figure 7:**
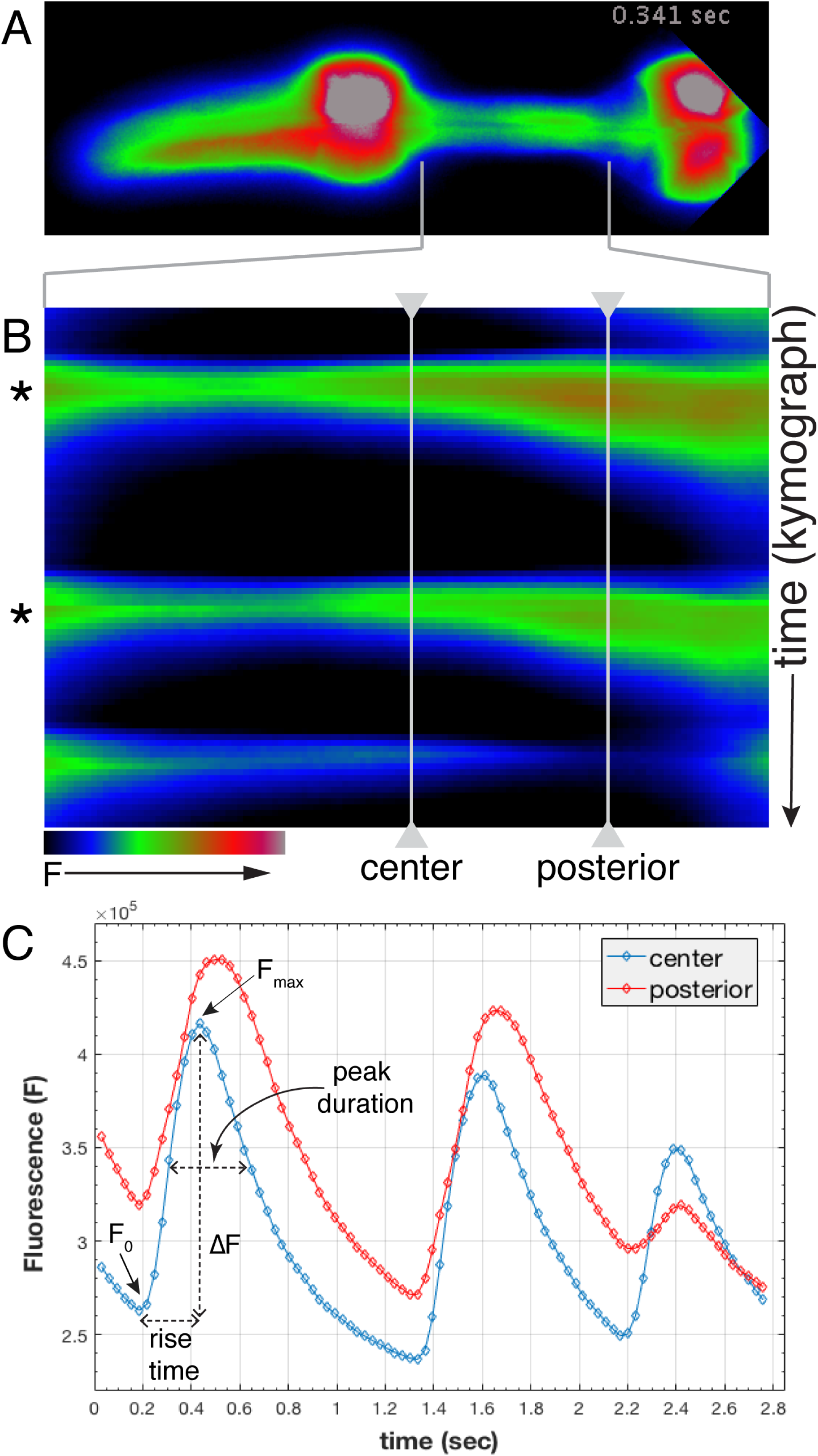
Characterization of GCaMP3 fluorescence in the pharyngeal isthmus. A) False colored fluorescence image of a wild-type animal expressing GCaMP3 in the pharyngeal muscles (anterior is left). B) Kymograph of maximum GCaMP3 fluorescence intensity in the isthmus region (indicated by brackets in A and B). Two pumps with peristalsis marked with asterisks are followed by a pump without a peristalsis. C) Fluorescence levels (F) in the center and posterior isthmus plotted vs time (sec) for the contractions in panel B. Time points for time-lapse imaging (circles) and example measurements for ∆F, peak duration and rise time are indicated.

To compare Ca^2+^ transients in the center and posterior isthmus in wild-type animals to those in various mutants, we quantified for each pump and peristalsis the normalized change in fluorescence (∆F/F_o_), the width of each peak at 50% maximum fluorescence (peak duration), and the time that it took for fluorescence to reach its maximum (rise time) (Table 2; Figure 7 C). In addition we calculated the time delay between the maximum increase in GCaMP3 fluorescence in the center and posterior isthmus (peak delay).

**Table 2:** Quantification of GCaMP3 fluorescence dynamics

| genotype | isthmus position | normalized $\Delta F$ | peak duration (ms±sem) | rise time (ms±sem) | peak delay (ms±sem) |
| --- | --- | --- | --- | --- | --- |
| wild type <sup>a</sup> | center | 52±2 | 329±7 | 256±5 | |
| | posterior | 40±2 | 376±10 | 279±6 | 33±4 |
| *eat-2(ok3528)* <sup>b</sup> | center | 78±3 | 598±30 | 432±9 | |
| | posterior | 63±3 | 666±39 | 426±12 | 67±7 |
| *gar-3(gk305)* <sup>c</sup> | center | 51±3 | 310±11 | 258±8 | |
| | posterior | 34±2 | 345±14 | 281±10 | 7±3 |
| *eat-2; gar-3* <sup>d</sup> | center | 57±2 | 414±12 | 315±6 | |
| | posterior | 36±2 | 485±17 | 346±8 | 26±4 |

<sup>a</sup> 15 wild type OK1020 young adults were recorded and an average of 130 pumps were analyzed.
<sup>b</sup> Six *eat-2(ok3528)* young adults were recorded and an average 56 pumps were analyzed. These animals exhibited increased normalized $\Delta F$ , peak duration and rise time in both the center and posterior isthmus ( $p < 10^{-7}$ ) and an increased peak delay ( $p < 10^{-4}$ ) compared to wild type.
<sup>c</sup> Five *gar-3(gk305)* young adults were recorded and an average of 31 pumps were analyzed. These animals exhibited a decreased peak delay ( $p < 10^{-5}$ ) compared to wild type.
<sup>d</sup> 12 *eat-2; gar-3* young adults were recorded and an average of 94 pumps were analyzed. These animals exhibited decreased normalized $\Delta F$ , peak duration, rise time, and peak delay compared to *eat-2(ok3528)* single mutants ( $p < 10^{-5}$ ).

We found changes in GCaMP3 fluorescence in the pharyngeal isthmus that paralleled many of the defects we observed in peristaltic contractions in various mutants (Figure 8; Table 2). *eat-2(ok3528)* mutants, which have prolonged peristalses, also exhibited significantly increased ∆F/F_o_, peak duration and rise time in both the center and posterior isthmus. In addition, the peak delay was increased 2-fold (Movie S8 & S9). In comparison, *gar-3(gk305)* mutants, which have normal duration of peristalses, exhibited ∆F/F_o_ and rise times that were similar to those of wild-type animals, with a small decrease in ∆F/F_o_ only in the posterior isthmus, as well as a small decrease in peak duration in both the center and posterior isthmus. However, the most striking change in these animals was a strongly decreased peak delay between the center and posterior isthmus. Finally, *eat-2(ok3528); gar-3(gk305)* double mutants, which suppress the prolonged peristalses observed in *eat-2(ok3528)* single mutants, exhibited decreases in ∆F/F_o_, peak duration, rise time, and peak delay compared to *eat-2(ok3528)* single mutants, although only ∆F/F_o_ was reduced to wild-type levels. Taken together, these results show that the level, duration, and rise time of Ca^2+^ increases in both the center and posterior isthmus best correlate with the duration of peristaltic contractions. Loss of *eat-2* increases these Ca^2+^ signals and produces prolonged peristalses that depend on wild-type *gar-3*.

**Figure 8:**
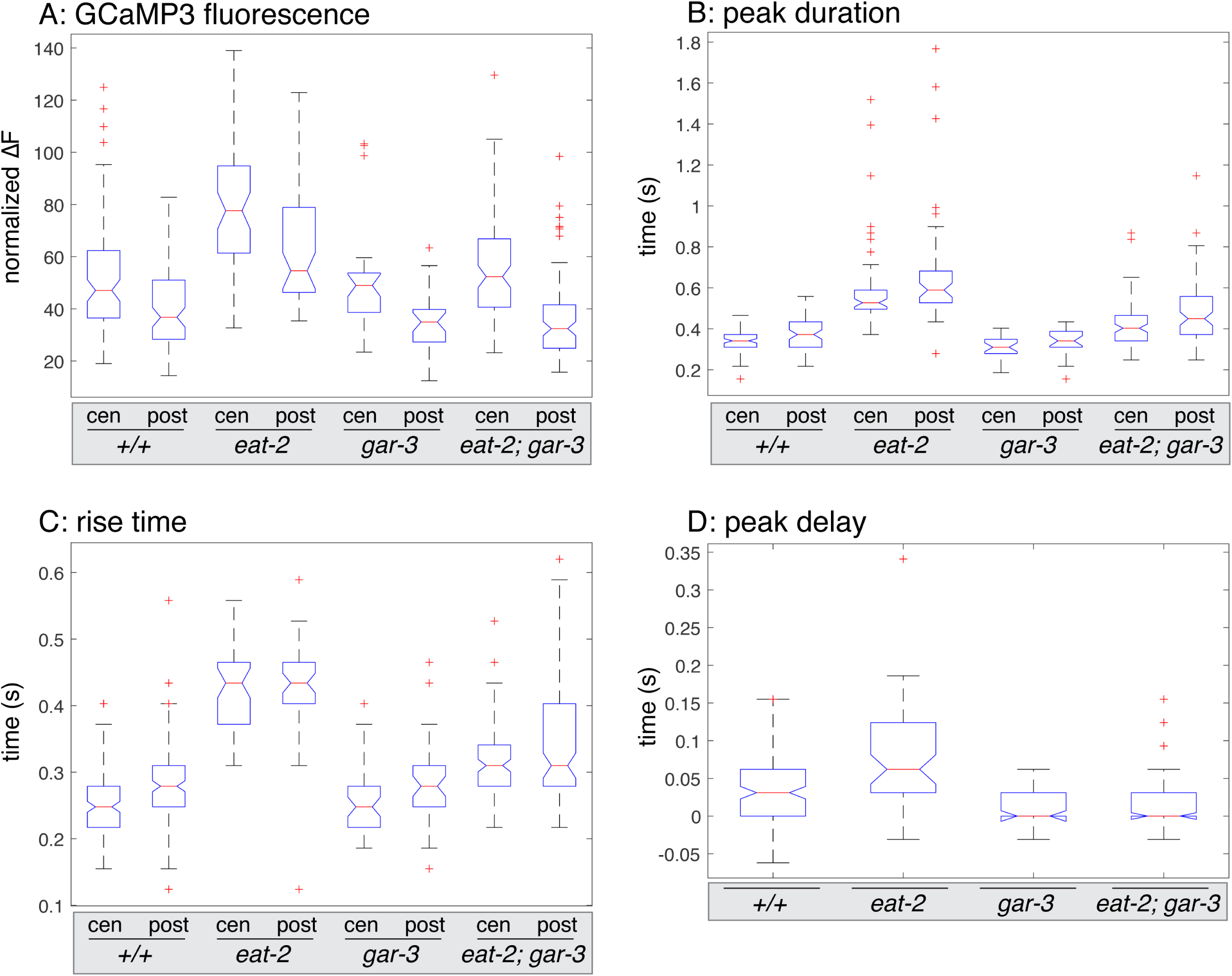
Quantification of GCaMP3 fluorescence in wild-type animals and mutants. Box and whisker plots comparing values measured from time-lapse imaging of GCaMP3 fluorescence during pumps followed by peristalses: (A) normalized GCaMP3 fluorescence levels (∆F/F_o_), (B) peak duration, (C) peak rise time, (D) peak delay between the center and posterior isthmus. Genotypes and measurements in the center (cen) and posterior (post) isthmus are indicated. The central bar (red) denotes the median value with a notch indicating the 95% confidence for this median, the box indicates the interquartile range (IQR, 25th to 75th percentile), and the whiskers indicating values within 1.5 times the IQR. Suspected outlier values are indicated as red '+'.

### *ace-3* mutants exhibit prolonged peristalses that depend on GAR-3

It is surprising that *eat-2* loss leads to a reduced pump rate, while at the same time hyperstimulating peristalsis and increased cytoplasmic Ca^2+^ concentration in the isthmus muscles. One hypothesis for these unexpected effects is that, in *eat-2* mutants, ACh released from MC spills over from the synapses and stimulates GAR-3 receptors located in the isthmus muscles. To examine this hypothesis, we characterized pharyngeal muscle contractions in *ace* mutants defective in acetylcholinesterases that normally hydrolyze ACh and have increased unbound ACh in the synapse (Johnson *et al.* 1988; Combes *et al.* 2000). *ace-3(dc2)* mutants exhibited significantly prolonged peristalses, which were similar to those of *eat-2* mutants (Table 1). In comparison, *ace-2(g72); ace-1(p1000)* double mutants exhibited only mildly prolonged peristalses. Both of these strains also exhibited a small decrease in the pump rate, but this was not significantly different from wild type, indicating that MC can effectively stimulate pumping in *ace* mutants.

To ask if the prolonged peristalses were suppressed by *gar-3* mutation, we examined *ace-3(dc2); gar-3(gk305)* double mutants and found that these animals exhibited peristalses that were similar to wild type (Table 1). Thus the prolonged peristalses in *ace-3* mutants depend on crosstalk with wild-type GAR-3 receptor similar to what we observed in *eat-2* mutants.

### *gar-3* suppresses the slow growth and extended life span of *eat-2* mutants

As *gar-3* mutation suppresses the peristalsis defects in *eat-2* mutants, we wanted to determine if this mutation also affects the persistent feeding defects in *eat-2* mutants. *eat-2* mutants exhibit slow larval growth and a prolonged adult life span, which result from dietary restriction (Avery 1993a; Lakowski and Hekimi 1998). We found that *gar-3* mutation at least partially suppressed both of these phenotypes.

While nearly all wild-type animals and *gar-3(gk305)* single mutants reached adulthood after 5 days at 20°C, *eat-2* mutants grew much more slowly, and, even after 6 days, only 30% had reached adulthood (Figure 9 A). *eat-2(ok3528); gar-3(gk305)* double mutants also grew slower than wild type and *gar-3(gk305)*, but they grew faster than *eat-2(ok3528)* single mutants, and 100% of these animals reached adulthood within 6 days.

**Figure 9:**
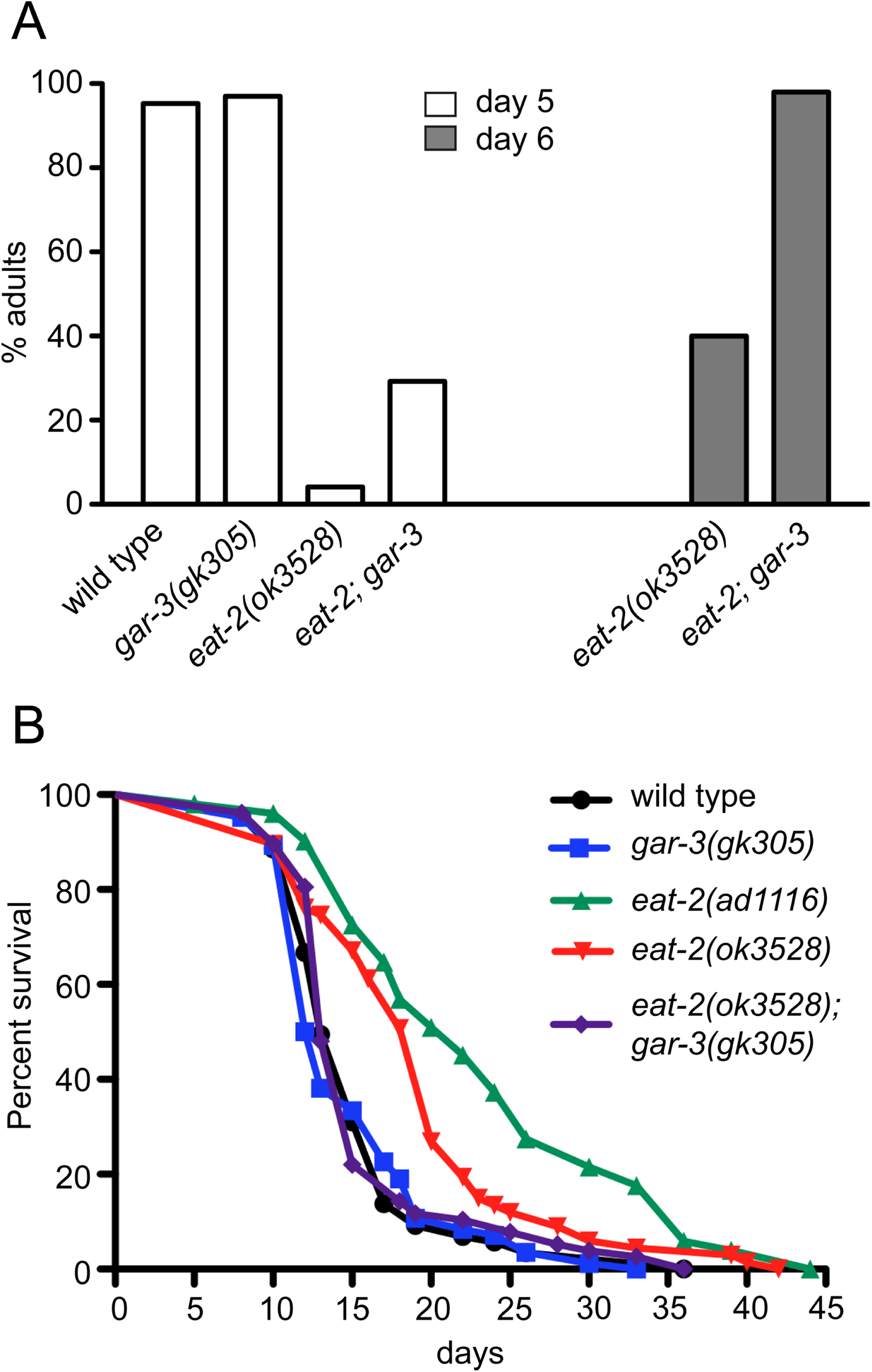
*gar-3* mutation suppresses the slow growth and prolonged life span of *eat-2* mutants. A) Bar graphs indicating the percent animals of the indicated genotypes reaching adulthood by day 5 or day 6 at 20°C (n = 100 for each genotype). B) Adult survival curves for animals of the indicated genotypes. Averages of representative triplicate assays performed on plates containing UV-killed *E. coli* as a food source. Nearly identical results were obtained in assays with living *E. coli*.

Likewise, *eat-2* mutants have previously been shown to exhibit a longer adult life span than wild type animals (Lakowski and Hekimi 1998), and we found that both *eat-2(ad1116)* and *eat-2(ok3528)* mutants exhibited significantly extended life spans (median adult survival 22 and 20 days, respectively) (Figure 9 B). In comparison, the life span of *gar-3(gk305); eat-2(ok3528)* double mutants was indistinguishable from that of wild-type animals and *gar-3(gk305)* single mutants [median adult survival: N2 (13 days), *gar-3(gk305)* (12.5 days), *gar-3(gk305); eat-2(ok3528)* (13 days)].

Thus *gar-3(gk305)* suppresses both the slow larval growth and extended life span phenotypes of *eat-2(ok3528)* mutants, and we suggest this suppression is the result of improved isthmus peristalsis and feeding.

## Discussion

In this work we show that crosstalk between the nAChR EAT-2 and the mAChR GAR-3 affects peristaltic muscle contractions in the isthmus of the *C. elegans* pharynx. Under normal circumstances, GAR-3 has a relatively minor role in these contractions, but in *eat-2* mutants, GAR-3 can stimulate increased cytoplasmic Ca^2+^ levels and peristalses that are prolonged compared to those of wild-type animals. We suggest this GAR-3 dependent stimulation of peristalsis results from ACh spillover from synapses between MC and the pharyngeal muscles in *eat-2* mutants, and consistent with this suggestion, acetylcholinesterase mutants similarly exhibit prolonged peristalses that are dependent on GAR-3. Crosstalk with GAR-3 contributes to the slow larval growth and extended life span phenotypes observed in *eat-2* mutants, indicating that this crosstalk has a long-term impact on *C. elegans* feeding.

### *eat-2* mutants exhibit different effects on pumping and peristalsis

*eat-2* mutants have a reduced pump rate, indicating that EAT-2 plays an excitatory role in pumping (Avery 1993b; Raizen *et al.* 1995). In comparison, *eat-2* mutants exhibit prolonged peristalses, indicating EAT-2 does not excite isthmus muscle contractions. Rather it is necessary for rapid relaxation. These peristalsis phenotypes are similar to previous observations demonstrating that *eat-2* mutation leads to prolonged depolarization and contraction of the terminal bulb muscles during pumping (Steger and Avery 2004).

The EAT-2 receptor is specifically localized at the sites where MC forms synapses near the junction of the pm4 and pm5 pharyngeal muscles, and it is necessary for MC to directly stimulate rapid pumping (Mckay *et al.* 2004). Because MC does not synapse on either the posterior isthmus or the terminal bulb (Albertson and Thomson 1976; Mckay *et al.* 2004), the prolonged contraction of these muscles is likely an indirect effect of loss of EAT-2 containing receptors.

### GAR-3 stimulates prolonged peristalsis in *eat-2* mutants

The prolonged peristalses in *eat-2* mutants depend on the GAR-3 receptor. *gar-3* mutation partially suppressed the prolonged peristaltic contraction of *eat-2* mutants, as well as the increased and prolonged Ca^2+^ transients in the isthmus muscles of these animals. This suppression demonstrates that GAR-3 receptor activation contributes to the *eat-2* mutant peristaltic phenotypes. Previous studies demonstrated that *gar-3* mutation similarly suppressed the prolonged contraction of the terminal bulb muscles in *eat-2* mutants (Steger and Avery 2004). In comparison, *gar-3* single mutants have relatively minor defects in peristalsis. The duration of peristalsis was similar to wild-type animals. However, these mutants exhibited a sharp decrease in the peak delay between Ca^2+^ transients in the center and posterior isthmus and a decreased ∆F/F_o_ in the posterior isthmus. Thus, in wild-type animals, GAR-3 contributes to the spatially and temporally dynamic changes in Ca^2+^ concentration in the isthmus muscles, but this did not produce recognizable changes in the duration of peristalsis. These observations demonstrate that GCaMP3 may be a more sensitive assay for muscle excitation.

In contrast to its effect on peristalsis, *gar-3* mutation did not suppress reduced pump rate observed in *eat-2* mutants (Steger and Avery 2004). Thus, the pumping and peristalsis defects in *eat-2* mutants arise via different mechanisms. Reduced pump rate results directly from loss of EAT-2 excitatory activity stimulated by the MC neuron, while the prolonged peristalsis results from excitation that depends on GAR-3.

### ACh spillover may produce prolonged peristalses in *eat-2* mutants

Extrasynaptic GAR-3 has previously been shown to respond to humoral ACh (Chan *et al.* 2013), and we suggest that in *eat-2* mutants ACh spillover from synapses between MC near the junction of the pm4 and pm5 muscles similarly activates GAR-3 receptors in pm5. Activation of the slow-acting, metabotropic GAR-3 stimulates increased and prolonged Ca^2+^ transients in the isthmus, which lead to prolonged peristalsis. GAR-3 is known to regulate Ca^2+^-dependent processes in the pharyngeal muscles, and increased GAR-3 signaling produces prolonged pharyngeal muscle contractions during pumping (Steger and Avery 2004). Because *gar-3* mutation only partially suppresses the prolonged peristalsis in *eat-2* mutant, other mechanisms must also contribute to the prolonged peristalsis in *eat-2* mutants. For example, voltage activated K^+^ channels such as EXP-2 might not be efficiently activated in *eat-2* mutants for rapid repolarization of the isthmus muscles (Davis *et al.* 1999; Shtonda and Avery 2005).

At cholinergic synapses, unbound ACh is normally degraded by acetylcholinesterases located at the synapse [reviewed in (Rotundo 2003)]. However, ACh released at mammalian NMJs can coactivate mAChRs to stimulate vasodilation in nearby blood vessels (Welsh and Segal 1997), although the physiological significance of this stimulation is uncertain (Hong and Kim 2017). ACh spillover has also been observed when quantal content is increased or when acetylcholinesterases are inhibited (Stanchev and Sargent 2011; Petrov *et al.* 2014). We have similarly observed that *C. elegans ace-3* mutants with reduced acetylcholinesterase activity produce prolonged peristalses that are dependent on GAR-3.

While MC morphology and synapse positioning is normal in *eat-*2 mutants, we cannot rule out that other defects in synapse formation or remodeling occur in these animals. Interestingly, loss of the gamma-subunit of the nAChR leads to a diffuse distribution of acetylcholinesterase clusters during development of mouse diaphragm muscles (Liu *et al.* 2008), and a similar defect in acetylcholinesterase localization could underlie signaling from the MC to the pharyngeal isthmus muscles in *eat-2* mutants.

Increased cytoplasmic Ca^2+^ levels in muscle cells have been observed in a number of human neuromuscular disorders, including myotonic dystrophies, Duchenne muscular dystrophy, as well as in *ColQ* and slow-channel myasthenic syndromes [reviewed in (Vallejo-Illarramendi *et al.* 2014; Engel *et al.* 2015)]. In particular, myotonic dystrophies are initially characterized by delayed relaxation of muscles after contraction, and this resembles the prolonged peristalsis that we have observed in *eat-2* and other mutants. Thus *eat-2* mutants may provide a new model to characterize the immediate effects of Ca^2+^ dysregulation in muscle cells.

## Acknowledgments

The authors are indebted to Paul Huber, Angenee Milton, Zhihua Li, Janet Richmond, Sreekanth Chalasani, Sebnem Ece Eksi, Ian Hope and many MCDB colleagues for reagents, strains and advice. This project was supported by the NIH/NIGMS (R01 GM82865), a UIC Campus Research Board (CRB) Pilot grant, and a UIC LAS Award for Faculty in the Natural Sciences. Some strains were provided by the CGC, which is funded by NIH Office of Research Infrastructure Programs (P40 OD010440).

## Supporting Information Legends

**Figure S1:**
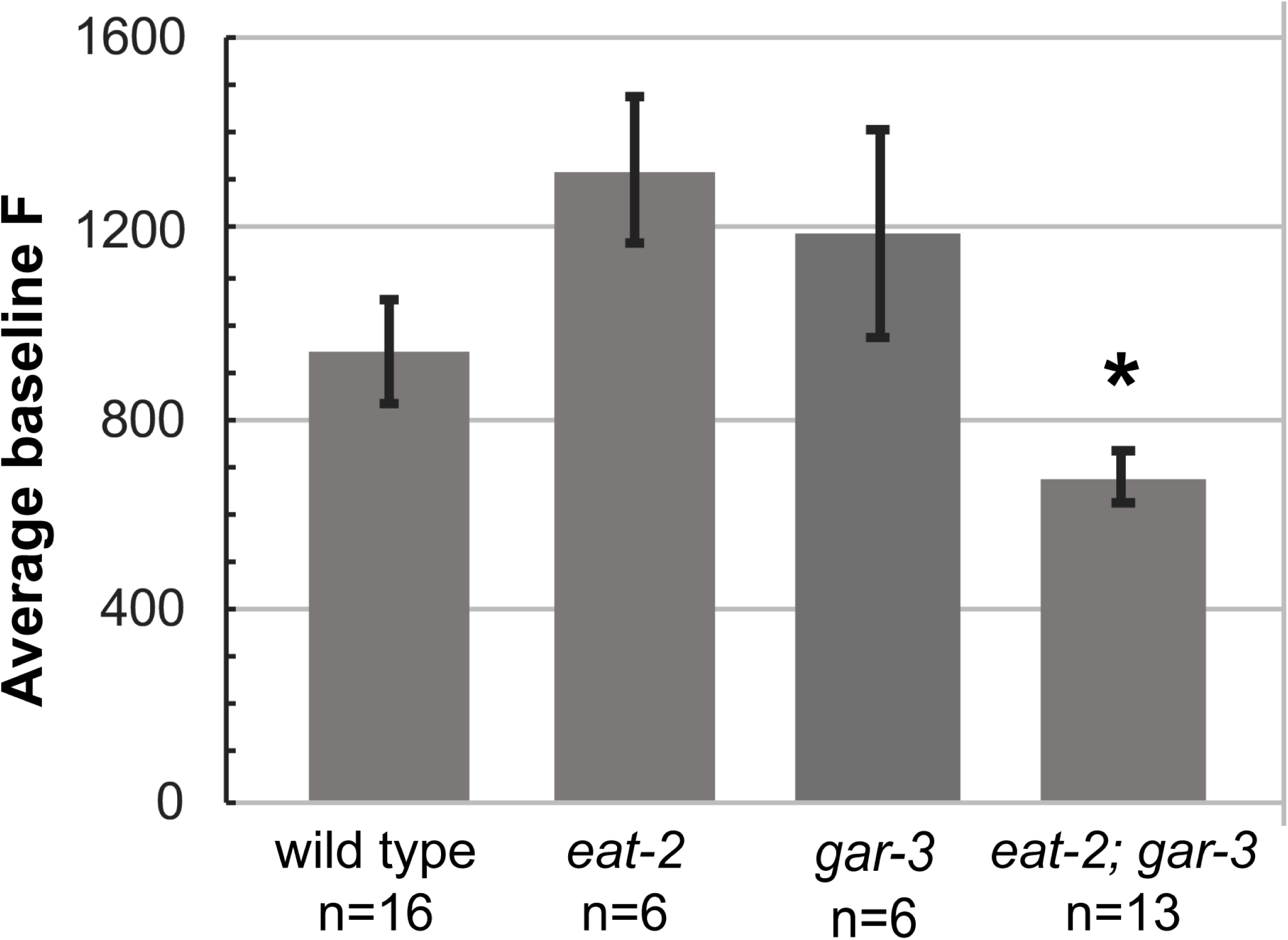
Average baseline GCaMP3 fluorescence intensity in the isthmus of adult animals of the indicated genotypes bearing the *myo-2^Prom^::GCaMP3* reporter. Images were acquired from animals under identical conditions and exposure times. Asterisk (*) indicates a significant difference from wild type values (p<0.05).

**Figure S2:**
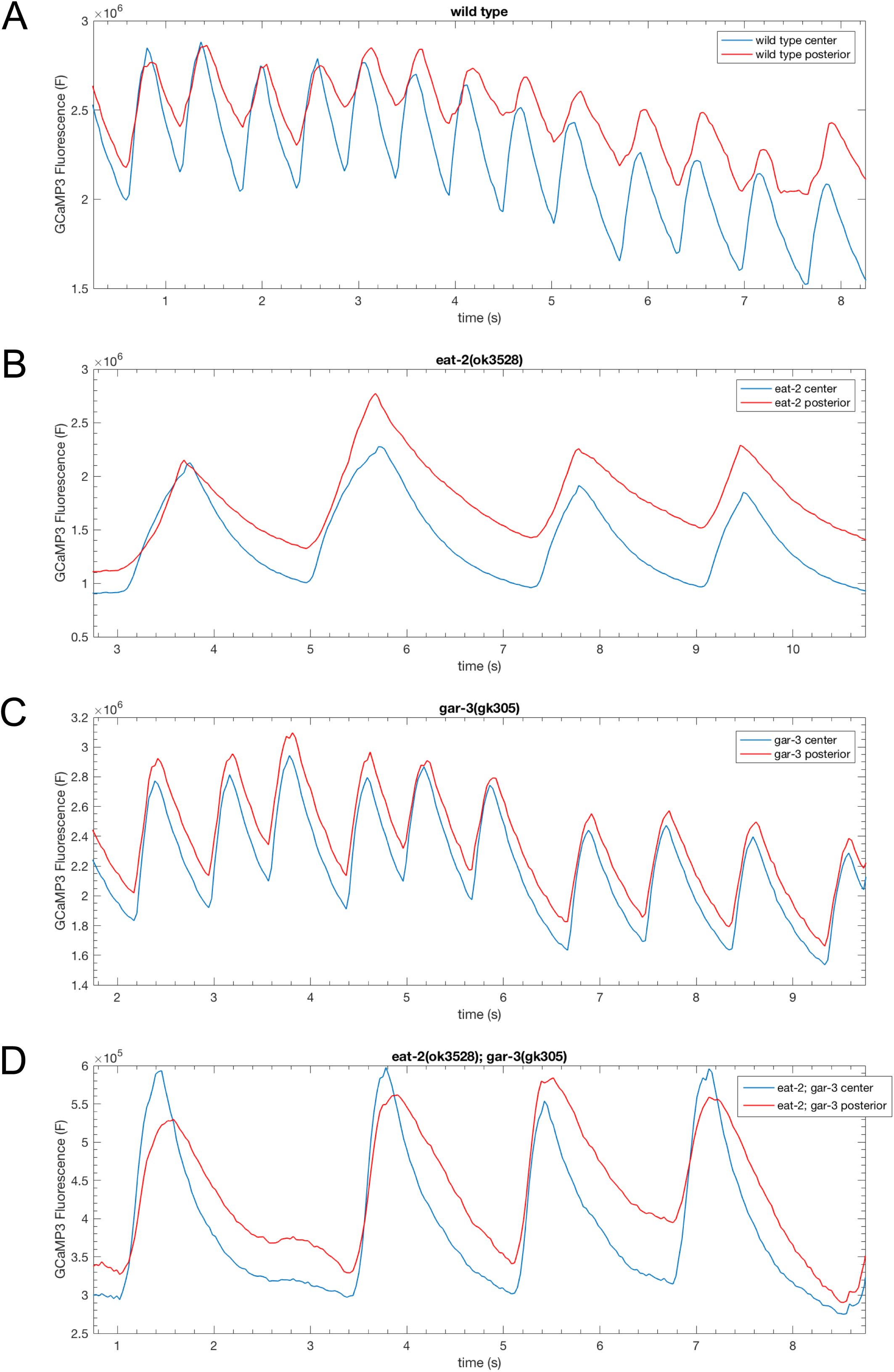
Eight second traces of fluorescence intensity (F) in the center (blue) and posterior (red) isthmus of representative animals of the indicated genotypes bearing the *cuIs36 myo-2^Prom^::GCaMP3* transgene. The data was extracted from (A) Table S14, (B) Table S29, (C) Table S36, and (D) Table S46 at the indicated time points.

**Movie S1:** DIC time-lapse movie of a wild-type L1 animal showing three pumps and one peristalsis after the second pump. Images were acquired at 25 frames/sec and are played back at 1/5 speed with the elapsed time shown in seconds.

**Movie S2:** DIC time-lapse movie of *cha-1(ok2253)* mutant lacking any pharyngeal muscle contractions. Images were acquired at 25 frames/sec and are played back at 1/5 speed with the elapsed time shown in seconds.

**Movie S3:** DIC time-lapse movie of *cha-1(ok2253)* mutant treated with 5mM nicotine showing two pumps, each followed by a peristalsis. Images were acquired at 25 frames/sec and are played back at 1/5 speed with the elapsed time shown in seconds.

**Movie S4:** DIC time-lapse movie of *cha-1(ok2253)* mutant treated with 5mM arecoline showing three pumps, each followed by a peristalsis. Images were acquired at 25 frames/sec and are played back at 1/5 speed with the elapsed time shown in seconds.

**Movie S5:** DIC time-lapse movie of *eat-2(ok3528)* L1 animal showing two pumps, each followed by a prolonged peristalsis. Images were acquired at 25 frames/sec and are played back at 1/5 speed with the elapsed time shown in seconds.

**Movie S6:** GCaMP3 time-lapse movie of a wild-type pharynx showing Ca^2+^ transients in the isthmus muscles during three pumps with first two followed by peristalsis. Images were acquired at 32 frames/sec and false colored (blue – low fluorescence, red – high fluorescence). Time-lapse is played back at 5 frames/sec.

**Movie S7:** Upper panel represents GCaMP3 time-lapse movie of a cropped isthmus from the same wild-type animal as shown in Sup. Movie 6. Bottom panel represents profile plot of GCaMP3 fluorescence intensity in the isthmus shown in the upper panel. GCaMP3 fluorescence is on the X-axis and distance in the isthmus is on Y-axis. Arrow shows wave like Ca^2+^ transients traveling through the posterior isthmus during peristalsis.

**Movie S8:** GCaMP3 time-lapse movie of *eat-2(ok3528)* pharynx showing Ca^2+^ transients in the isthmus muscles during one pump followed by a prolonged peristalsis. Images were acquired at 32 frames/sec and false colored (blue – low fluorescence, red – high fluorescence). Time-lapse is played back at 5 frames/sec.

**Movie S9:** Upper panel represents GCaMP3 time-lapse movie of a cropped isthmus from the same *eat-2(ok3528)* animal as shown in Sup. Movie 7. Bottom panel represents profile plot of GCaMP3 fluorescence intensity in the isthmus shown in the upper panel. GCaMP3 fluorescence is on the X-axis and distance in the isthmus is on Y-axis.

**Tables S2-S13: Time-lapse of pharyngeal muscle contractions**

Microsoft Excel files containing raw data extracted from time-lapse DIC images of pharyngeal muscle contractions quantified in Table 1. Each Excel file contains multiple tabs, which refer to individual animals, and each tab contains seven columns: frame (frame number), time (time in seconds), PC (procorpus), TB (terminal bulb), ant isth (anterior isthmus), mid isth (middle isthmus), post isth (posterior isthmus). Yellow shaded cells indicate times when the pharyngeal lumen was open (pharyngeal muscle contraction), while unshaded cells indicate frames when the pharyngeal lumen was closed (pharyngeal muscle relaxed). **Table S2** wild type, **Table S3** wild type + arecoline, **Table S4** *gar-3(gk305)*, **Table S5** *gar-3(gk305)* + arecoline, **Table S6** *eat-2(ad465)*, **Table S7** *eat-2(ad1116)*, **Table S8** *eat-2(ok3528)*, **Table S9** *eat-2(ok3528); gar-3(gk305)*, **Table S10** *eat-18(ad820)*, **Table S11** *ace-2(g72); ace-1(p1000)*, **Table S12** *ace-3(dc2)*, and **Table S13** *ace-3(dc2); gar-3(gk305)*.

**Tables S14-S52: Time-lapse of GCaMP3 fluorescence**

Microsoft Excel files containing GCaMP3 fluorescence measurements in the pharyngeal isthmus extracted from time-lapse fluorescence image quantified in Table 2. For each genotype there are multiple Excel files referring to individual animals, and each Excel file has two or more tabs referring to GCaMP3 fluorescence measurements in the central (Central) and posterior (Posterior) isthmus. Some Excel files have more than two tabs which reflect measurements for separate time periods of the same animal (frame number is indicated in parenthesis). Each tab has three columns: frame (frame number), time, sec (time in seconds) and F total (GCaMP3 total fluorescence). **Tables S14-S28** wild type, **Tables S29-S34** *eat-2(ok3528)*, **Tables S35-S40** *gar-3(gk305)*, and **Tables S41-S52** *eat-2(ok3528); gar-3(gk503)*.

**Table S1:** Quantification of pharyngeal muscle contractions in *acr* mutants

| genotype | pump rate<br>(pumps/min ±<br>sem) | peristalsis rate<br>(peristalses/min ±<br>sem) | % pumps<br>followed by<br>peristalsis | duration of<br>isthmus<br>peristalsis<br>(ms ± sem) |
| --- | --- | --- | --- | --- |
| wild type <sup>a</sup> | 218 ± 11 | 31 ± 3 | 14% | 113 ± 4 |
| *acr-6(ok3117)* <sup>b</sup> | 221 ± 14 | 12 ± 4 | 6% | 115 ± 13 |
| *acr-7(tm863)* <sup>c</sup> | 198 ± 3 | 14 ± 4 | 7% | 119 ± 7 |
| *acr-10(ok3118)* <sup>d</sup> | 199 ± 29 | 20 ± 6 | 12% | 96 ± 3 |
| *acr-14(ok1155)* <sup>e</sup> | 199 ± 29 | 21 ± 6 | 12% | 96 ± 4 |
| *acr-16(ok789)* <sup>f</sup> | 184 ± 9 | 20 ± 2 | 12% | 118 ± 7 |

<sup>a</sup> 17 N2 L1s were recorded for 19-20 s and a total of 1194 pumps were analyzed.
<sup>b</sup> Six *acr-6* L1s were recorded for 19-20 s and a total of 428 pumps were analyzed.
<sup>c</sup> Six *acr-7* L1s were recorded for 19-20 s and a total of 392 pumps were analyzed.
<sup>d</sup> 21 *acr-10* L1s were recorded for 20-21 s and a total of 1190 pumps were analyzed.
<sup>e</sup> Five *acr-14* L1s were recorded for 19-20 s and a total of 325 pumps were analyzed.
<sup>f</sup> Six *acr-16* L1s were recorded for 19-20 s and a total of 364 pumps were analyzed.

